# p53 is required for female germline stem cell maintenance in *P*-element hybrid dysgenesis

**DOI:** 10.1101/204800

**Authors:** Sadia Tasnim, Erin S. Kelleher

**Affiliations:** Department of Biochemistry and Molecular Biology, Institute for Translational The University of Texas Medical Branch, Galveston, TX 77555; Department of Biology and Biochemistry, University of Houston, Houston, TX 77204

## Abstract

Hybrid dysgenesis is a sterility syndrome resulting from the mobilization of certain transposable elements in the *Drosophila* germline. Particularly extreme is the hybrid dysgenesis syndrome caused by *P*-element DNA transposons, in which dysgenic female ovaries often contain few or no germline cells. Those offspring that are produced from dysgenic germlines exhibit high rates of *de novo* mutation and recombination, implicating transposition-associated DNA damage as the cause of germline loss. However, how this loss occurs, in terms of the particular cellular response that is triggered (cell cycle arrest, senescence, or cell death) remains poorly understood. We demonstrate that two components of the DNA damage response, Checkpoint kinase 2 and its downstream target p53, are determine the frequency of ovarian atrophy that is associated with *P*-element hybrid dysgenesis. We further show that p53 is strongly induced in the germline stem cells (GSCs) of dysgenic females, and is required for their maintenance. Our observations support the critical role for p53 in conferring tolerance of transposable element activity in stem cells.

## INTRODUCTION

Transposable elements (TEs) are mobile genetic parasites that ensure their spread through host populations by replicating in germline cells. This selfish replication not only harms the host by producing deleterious mutations that are transmitted to offspring (Dupuy et al., 2001; Spradling et al., 1999), it can also decrease host fertility by disrupting gametogenesis (Josefsson et al., 2006; Orsi et al., 2010; Schaefer et al., 1979). This is particularly clear with hybrid dysgenesis, a sterility syndrome of *Drosophila* caused by the unregulated transposition of certain TE families in germline cells (Bingham et al., 1982; Bucheton et al., 1984; Evgen’ev et al., 1997; Hill et al., 2016; Rubin et al., 1982; Yannopoulos et al., 1987). The hybrid dysgenesis syndrome imposed by *P*-element DNA transposons is particularly dramatic, causing females to have small atrophied ovaries that contain few or no germline cells (Dorogova et al., 2017; Schaefer et al., 1979). While the DNA damage associated with *P*-element transposition is thought to precipitate germline loss (Khurana et al., 2011), the cellular mechanisms that are responsible remain poorly understood (Dorogova et al., 2017; reviewed in Malone et al., 2015).

The DNA damage response (DDR) is a network of cellular pathways that detects and integrates DNA damage signals and determines cellular responses. In the *Drosophila* female germline, DNA damage is known to induce both cell-cycle arrest and apoptosis (Hassel et al., 2013; Shim et al., 2014), which could explain the absence of developing egg chambers in the presence of rampant transposition. In particular, the DDR inducer Checkpoint kinase 2 (Chk2) and its downstream effector p53 play key roles in determining the fate of damaged germline genomes. In early oogenesis, among germline stem cells (GSCs) and their mitotically-dividing daughter cystoblasts (CBs), these proteins have opposing functions. *mnk/loki*, which encodes Chk2, is required to trigger cell cycle arrest in response to DNA damage, while *p53* is required to ensure cell-cycle re-rentry (Ma et al., 2017; Shim et al., 2014; Wylie et al., 2014). By contrast, in early meiotic cells, *p53* acts downstream of *mnk/loki* to trigger apoptosis (Shim et al., 2014). Chk2 activation can also induce meiotic arrest in response to unrepaired double stranded breaks in a *p53*-independent manner (Abdu et al., 2002).

Although DDR has not been directly linked to dysgenic ovarian atrophy, two lines of evidence implicate it as a cause of germline loss. First, when sterility is incomplete, egg chambers produced by *P*-element dysgenic females often show morphological defects associated with Chk2-induced meiotic arrest (Khurana et al., 2011). Second, increased cell-death is observed in the early meiotic stages of dysgenic female germlines (Dorogova et al., 2017), the precise developmental time-point at which *mnk/loki* and *p53*-dependent apoptosis is known to result from DNA damage (Hassel et al., 2013; Shim et al., 2014).

In this study, we directly interrogate the role of the DDR inducers Chk1 and Chk2, and their downstream effector p53, in producing dysgenic ovarian atrophy. We demonstrate that *mnk/loki* and *p53* are strong genetic modifiers of hybrid dysgenesis, consistent with their roles in determining cellular response to DNA damage. We further reveal that p53 is induced in dysgenic GSCs and CBs, and is required for GSC maintenance in the dysgenic germline. Our results implicate DDR as a key determinant germline cell fate in the presence of *P*-element activity, and highlight the role of p53 in conferring GSC tolerance of DNA damage.

## MATERIALS and METHODS

***Drosophila stocks.** p53[5A-1-4], p53[11-1B-1]* (Rong et al., 2002) and Harwich (Kidwell et al., 1977) were obtained from the Bloomington stock center. *grp[209], grp[Z5170],* and *loki[30]* were generously provided by Jeff Sekelsky (LaRocque et al., 2007). *mnk[6006]* was generously provided by Bill Theurkauf. Canton-S was generously provided by Richard Meisel. *ep(2)1024/cyo;Hsp70-Gal4* was generously provided by Haifan Lin (Cox et al., 2000). Two p53 biosensors, *rpr-GFP-cyt* and *rpr-GFP-nuc*, were generously provided by John Abrams.

To extract the loss of function mutations into a Harwich genetic background, we first extracted 2^nd^ (*Cyo*) or 3^rd^ (*TM6*) balancer chromosomes into Harwich by crossing Harwich females to the multiply marked stock *w*; Sp/Cyo; sens[Ly-1]/TM6 Sb Tb* (Bloomington Stock number 42294), and selecting for balancer *(Cyo/+; TM6/+)* males, which were backcrossed to Harwich. The resulting marked stock was used for extraction of loss of function chromosomes. The final genotypes were: *H;loki[30]/Cyo;H, H;grp[209]/Cyo;H, H;grp[Z5170]/Cyo;H, H;H; p53[5A-1-4]/TM3 or TM6*, and *H;H; p53[11-1B-1]/TM3 or TM6*, where H indicates a Harwich chromosome. We did not extract the *mnk[6006]* allele into the Harwich background, because this allele is caused by a *P*-element insertion (Brodsky et al., 2004), and could therefore be unstable in the presence of P-transposase encoded by Harwich chromosomes.

**Crosses.** To evaluate the effects of DDR loss-of-function mutations on dysgenic ovarian atrophy we compared F1 offspring of crosses between M strain females that lack *P*-elements or maternally transmitted *P*-element regulation, and P strain males that contain active genomic *P*-elements that can induce hybrid dysgenesis (reviewed in Kelleher, 2016). To detect dominant effects, we crossed heterozygous females (DDR mutant/balancer, M background) to Harwich males. To evaluate the recessive effects we crossed heterozygous females (DDR mutant/balancer, M background) in to heterozygous (DDR mutant/balancer) Harwich (P) males. These crosses were performed and offspring were reared at 25°C, as we have previously observed that this temperature is optimal for revealing variation in the severity of dysgenesis (Srivastav and Kelleher, 2017).

To evaluate the effect of Piwi overexpression in somatic tissues, *ep(2)1024/cyo;Hsp70-Gal4* were crossed to Harwich and Canton-S males at 25°C. F1 female offspring were reared at 25°C, but exposed to a 1hr heat-shock (37°C) every 12 hours for 3 days (Cox et al., 2000).

**Ovarian atrophy.** 3-6 day-old female offspring were squashed under a coverslip with food-dye, to detect the presence of stage 14 egg chambers (Srivastav and Kelleher, 2017).

**Immunolabelling.** Ovaries were dissected from 3–6 day-old female offspring, and prepared according to the method of McKearin and Ohlstein (1995). Primary antibody concentrations were as follows: anti-GFP 1:1000, anti-Hts 1B1 1:4 (DSHB, Zaccai and Lipshitz, 1996), anti-Vasa 1:40. Secondary antibody concentrations were 1:500.

**Microscopy.** Ovaries were visualized with a SP8 Upright Confocal DM6000 CFS (Leica) Microscope outfitted with a 60X oil immersion lens. Images were gathered using an EM-CCD camera (Hamamatsu) and the LAS AF software (Leica).

## RESULTS

**Chk2 and p53 are strong modifiers of hybrid dysgenesis.** Dysgenic ovarian atrophy occurs among F1 offspring of crosses between M strain females who lack genomic *P*-elements and maternally transmitted *P*-element regulation, and P strain that carry a large number of active *P*-elements (Bingham et al., 1982; reviewed in Kelleher, 2016). We first considered the role of the DDR in dysgenic germline loss by determining whether loss-of-function alleles in DDR components are associated with an altered frequency of the ovarian atrophy phenotype. In addition to *mnk/loki* and *p53*, we also examined alleles of *grapes*, which encode the DDR inducer Checkpoint kinase 1 (Chk1). *grapes* works synergistically with *mnk/loki* in mitotic arrests in the female germline (Shim et al., 2014).

All the mutant alleles we examined were generated and maintained in M backgrounds (Bellen et al., 2004; Koundakjian et al., 2004; Rong et al., 2002; Rørth, 1996). We therefore looked for dominant effects of DDR alleles by examining the F1 offspring of heterozygous *(mutant/balancer*, M strain) females crossed to males of the reference P strain Harwich (Kidwell et al., 1977). For two separate alleles of *grapes*, a single allele of *mnk/loki*, and a single allele of *p53 (p53[5A-1-4])* we detected no dominant effects, as heterozygous F1 offspring *(mutant/+)* did not differ from their wild-type siblings *(balancer/+*, Figure 1A). A second *p53* allele *(p53[11-1B-1])* was associated with a marginally significant increase in dysgenic ovarian atrophy (p = 0.06), suggesting it is a mild dominant enhancer of ovarian atrophy.

To look for recessive effects, we crossed M strain female heterozygotes (*mutant/balancer*) to P strain male heterozygotes (*mutant/balancer*), from stocks we generated by extracting the mutant and balancer chromosomes into the Harwich background. We observed strong recessive effects of *mnk/loki* alleles on the frequency of dysgenic ovariation atrophy (Figure 1B). Transheterozygous *mnk/loki* offspring exhibited a dramatic 80% reduction in their incidence of ovarian atrophy when compared to their heterozygous (mutant/balancer) siblings. *mnk/loki* homozygotes could not be examined due to low viability *(loki[30])*, or instability of the causative insertion in the Harwich background *(mnk[6006]).grapes* alleles were weaker repressors than *mnk/loki*, with only one allele *(grp[Z5170])* showing significantly reduced atophy in homozygotes (30%) and transheterozygoes (39%). In contrast to the *mnk/loki* and *grapes*, homozygous and transheterozygous combinations of *p53* alleles enhanced ovarian atrophy by 13–22% when compared to heterozygous controls. The relative increase in ovarian atrophy is misleadingly modest, because it reflects almost complete sterility (99–100%) among dysgenic *p53* homozygotes and transheterozygotes.

Opposing effects of *mnk/loki* and *p53* mutations are consistent with the action of their gene products in GSCs, in which Chk2 triggers arrest in response to DNA damage, and p53 is required for cell-cycle re-entry (Ma et al., 2017; Wylie et al., 2014). If GSCs are arrested in atrophied ovaries, stimulating GSC division could restore gametogenesis. Somatic overexpression of *piwi* is known to non-autonomously stimulate GSC division (Cox et al., 2000). We therefore took advantage of a UASt-containing EP insertion in the 5’ piwi-UTR, which drives ectopic expression only in somatic cells when bound by Gal4 (Cox et al., 2000; Rørth, 1998). We crossed *ep(2)1024/Cyo;hsGal4* (M) females to Harwich (P) and Canton-S (M) males, and compared both classes of offspring after three days of heat-shock treatment. Consistent with GSC arrest, *piwi* overexpression *(ep(2)1024/hsGal4)* was associated with significantly decreased ovarian atrophy in dysgenic crosses involving Harwich males, but not in non-dysgenic crosses involving Canton-S males (Figure 1C). Because the *P{hsGal4}* and EP insertions are unstable in the dysgenic germline due the presence of P-transposase, the mild effect we observed likely underestimates the full impact of *piwi* overexpression. Taken together, our genetic analyses strongly implicate DDR in GSCs and CBs as an important determinant of *P*-element dysgenic ovarian atrophy.

**Figure 1.**
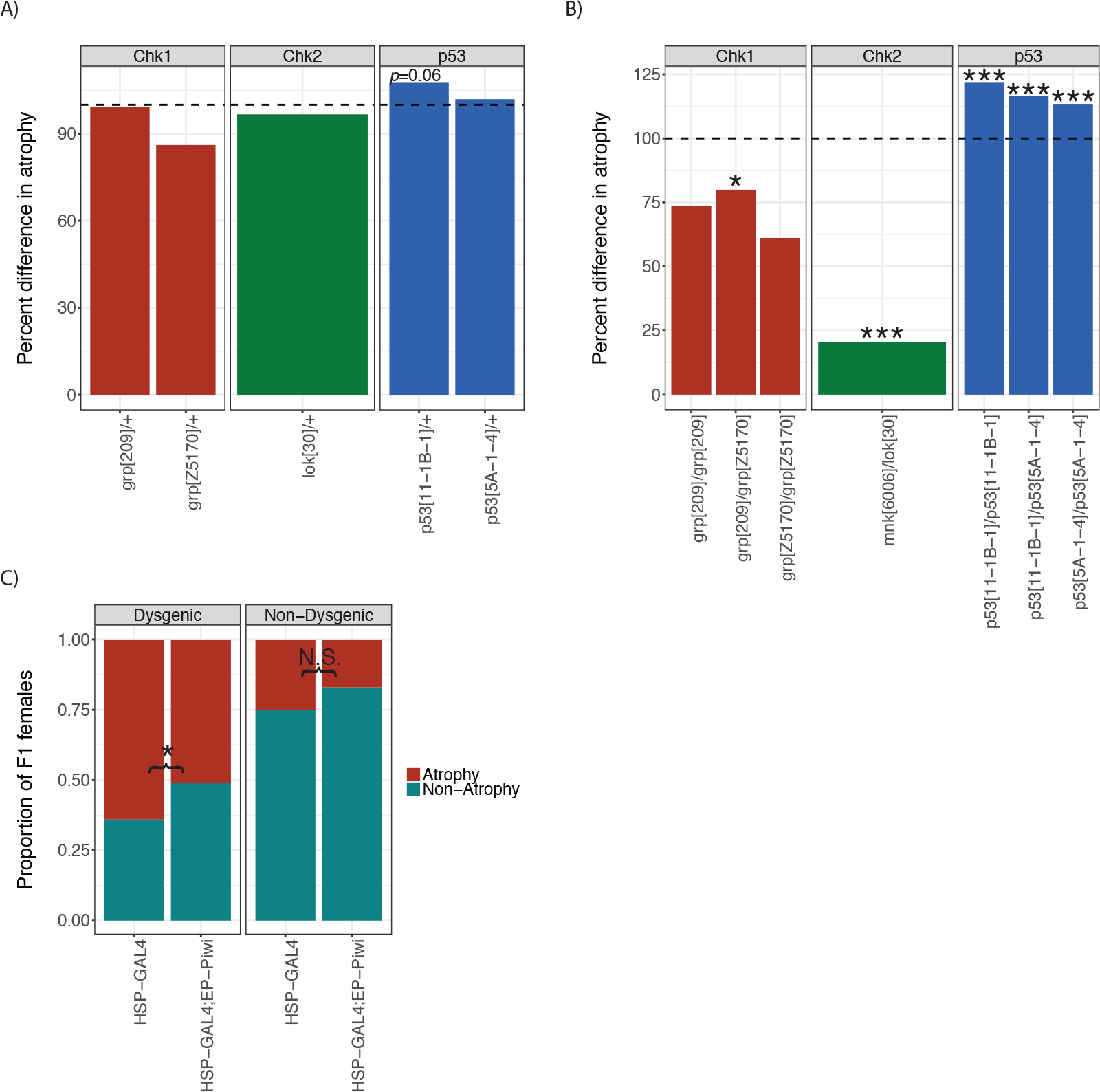
DDR components and regulators of GSC division are genetic modifiers of hybrid dysgenesis. The (A) dominant and (B) recessive effects of loss-of-function alleles for *grp, lok/mnk*, and *p53* on the incidence of F1 ovarian atrophy are evaluated. For each genotype, the frequency of ovarian atrophy is indicated as a percentage, relative to their homozygous wild type (A) and heterozygous (B) siblings from the same dysgenic cross. C) The proportion of offspring that exhibit ovarian atrophy is compared in the presence and absence of somatic *piwi* overexpression for dysgenic and non-dysgenic crosses. The statistical significance of the effect of each allele or genotype was ascertained by a *X*^*2*^ or Fisher’s exact test, and is indicated by the following notation. * denotes *p* ≥ 0.05, ** denotes *p* ≥ 0.01, *** denotes *p* ≥ 0.001, N.S. denotes non-significant.

**Nuclear p53 is induced in dysgenic GSCs.** p53 plays diverse roles in integrating DDR signals into cellular responses, both through its canonical role as a transcriptional regulator, and though interaction with both nuclear and cytoplasmic co-factors (reviewed in Green and Kroemer, 2009; Kastenhuber and Lowe, 2017). In the *Drosophila* germline, transcriptional activation by nuclear p53 occurs in response to DNA damage only in GSCs and CBs (Wylie et al., 2014). Given our observations that p53 is required for oogenesis in dysgenic females (Figure 1B), we wondered if p53 transcriptional activation is induced in these cells. To address this question, we took advantage of a previously developed p53 biosensor, in which GFP expression is controlled by regulatory sequences of the p53 target *reaper* (Lu et al., 2010). When p53 is present in the nucleus, GFP is produces. To compare p53 transcriptional activation in the presence and absence of *P*-element dysgenesis, we crossed females bearing two different p53 biosensors, encoding either nuclear or cytoplasmic GFP, to either Harwich (P) or Canton-S (M) strain males.

Consistent with the occurrence of DNA damage, we frequently observed induction of both biosensors in the GSCs or CBs of F1 females from dysgenic crosses involving Harwich males, but never in the non-dysgenic crosses involving Canton-S males (Figure 2A-C). This phenotype was incompletely penetrant, with 48% of F1 dysgenic germaria exhibiting GFP positive cells, as compared to 0% of non-dysgenic females. Incomplete penetrance of p53 induction could suggest that DNA damage associated with hybrid dysgenesis varies from cell-to-cell, but could also reflect stochastic excision of the biosensor transgene, which would be unstable in a *P-*dysgenic germline. Regardless, our observations suggests that p53 transcriptional regulation is induced by *P*-element activity in mitotically dividing GSCs and CBs, which is consistent evidence that *P*-elements often transpose in mitotic germline cells (Engels et al., 1990).

**Figure 2.**
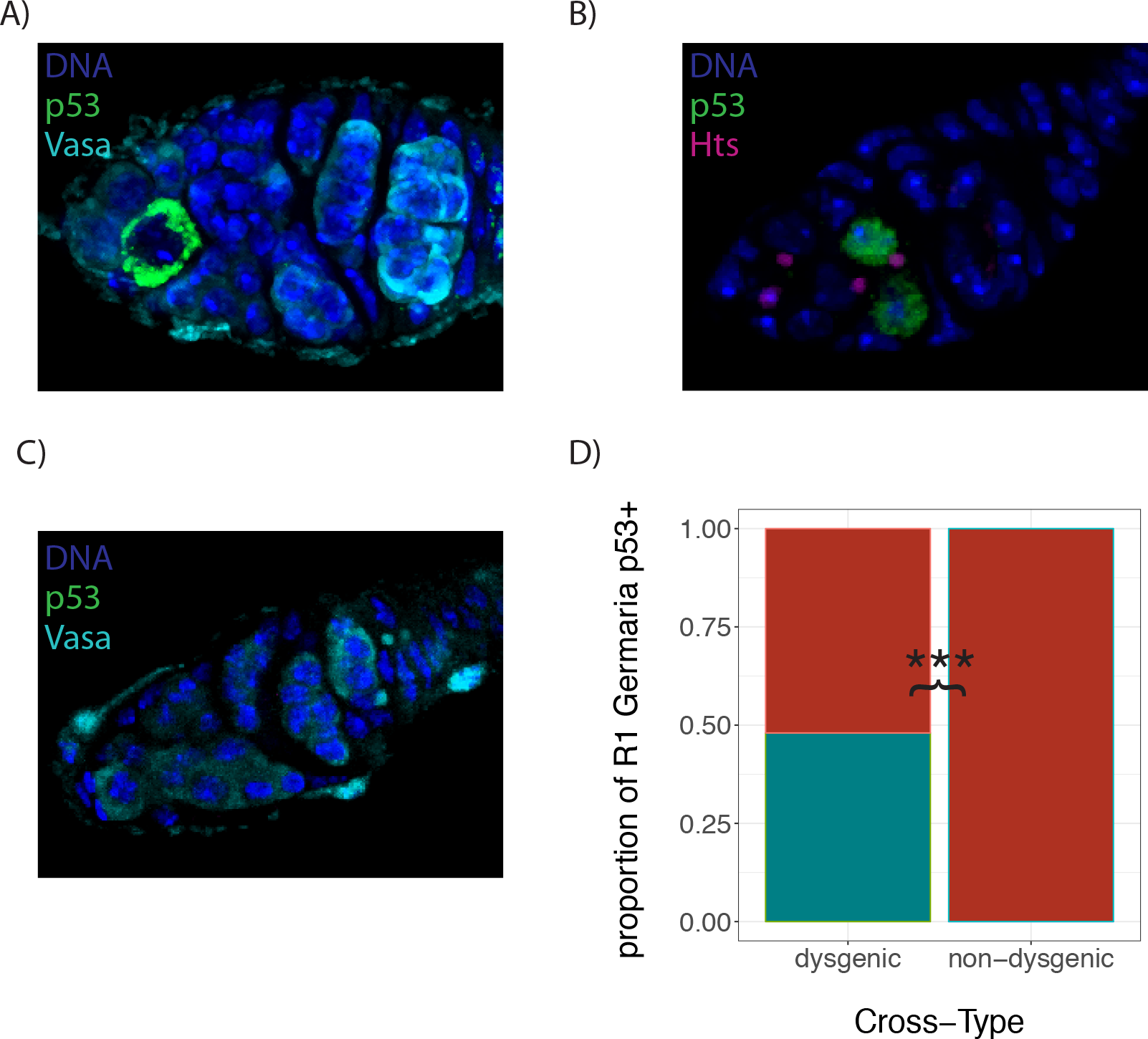
p53 is induced in dysgenic germaria. (A and B) Representative images of *rpr*-GFP expression in GSCs or CBs of dysgenic germaria, the ovarian structure where GSC reside and mitotic division occur (reviewed in Spradling, 1993). C) *rpr*-GFP is not induced in non-dysgenic germaria. All germaria are co-stained with DAPI to visualize nuclei, and either the germline marker Vasa (A and C), or Hts-1B1 (B), which identifies the circular spectrosomes of GSCs and CBs. D) The frequency of germaria with GFP positive cells in region 1, where GSCs and CBs occur, is compared between the dysgenic female offspring of Harwich fathers, and non-dysgenic offspring of Canton-S fathers. Statistical significance was ascertained by a Fisher’s exact test. *** denotes *p* ≥ 0.001

**p53 is required for GSC maintenance in dysgenic germaria.** Finally, we considered the role of p53 facilitating oogenesis in dysgenic females. We first examined the gross morphology of F1 female ovaries, to look for evidence of earlier-stage non-chorionated egg chambers, which would not be detected by the squash prep we employed for Figure 1. However, consistent with a requirement for *p53* at the earliest stages of oogenesis, dysgenic *p53* transheterozygotes overwhelmingly lacked any evidence of female germline development; with 55 of 56 females we dissected exhibiting two completely atrophied ovaries. (Figure 3A, D) In contrast, 14 out of 21 heterozygous females exhibited germline development in one or both ovaries (Figure 3B-D).

**Figure 3.**
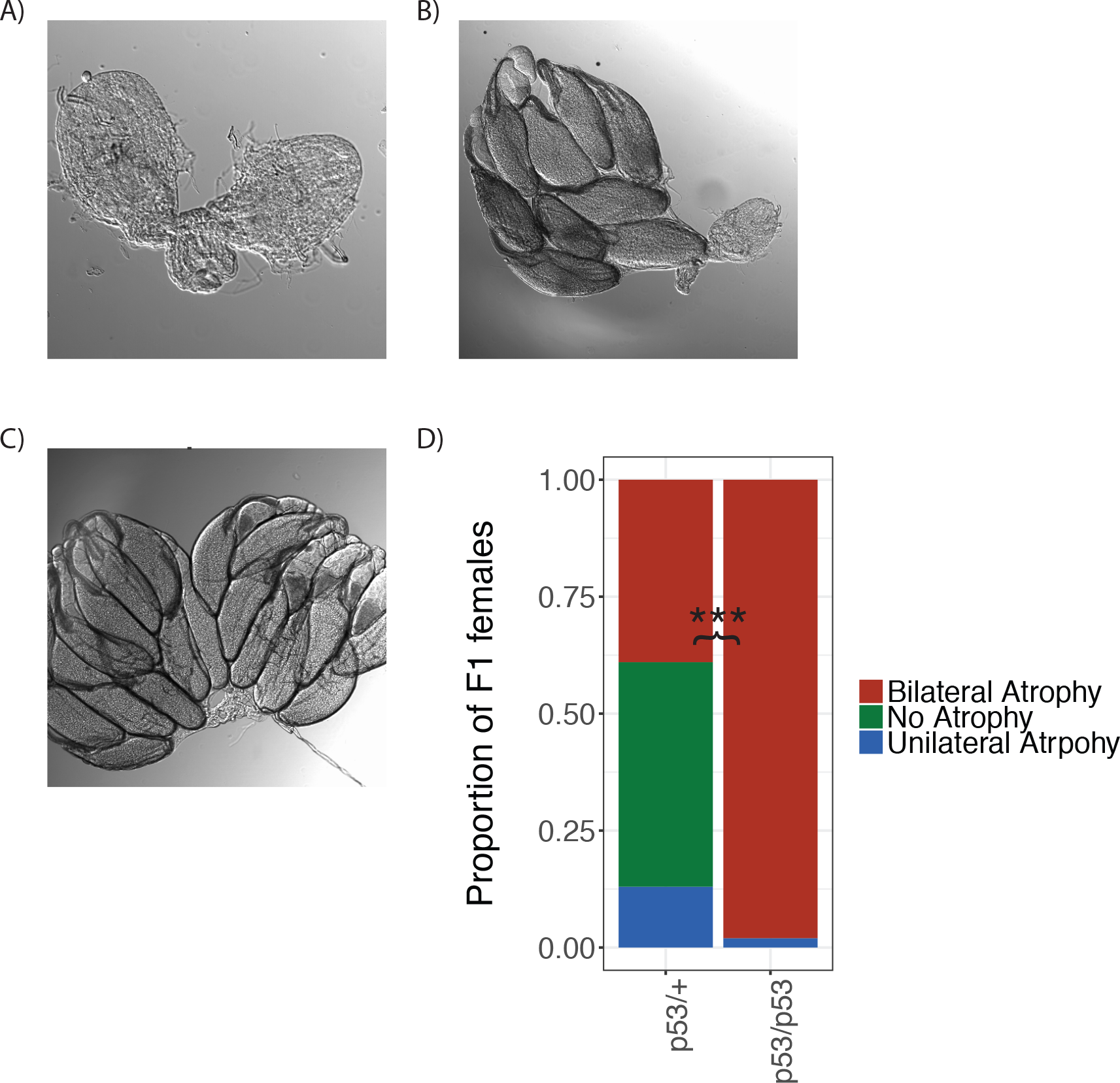
p53 mutants exhibit bilateral ovarian atrophy. Representative images if A) bilateral and B) unilateral ovarian atrophy are presented, as a well as pair of non-atrophied ovaries. D) The frequency of ovarian phenotypes is compared between *p53* heterozygotes (*p53[5A-1-4]*/+) and transheterozygotes (*p53[5A-1-4]/p53[11-1B-1]*). Statistical significance was assessed by a 2x2 *X*^*2*^ test comparing the frequency of bilateral atrophy, to that of unilateral and no-atrophy combined, between the two genotypes. *** denotes *p* ≥ 0.001

If *p53* is required only for cell cycle re-entry in damaged GSCs and CBs (Wylie et al., 2014), then dysgenic *p53* transheterozybotes should exhibit a very early pre-meiotic arrest in oogenesis. However, a recent study suggests that in mutants that are defective in TE regulation, p53 is also required for GSC maintenance (Ma et al., 2017). Consistent with a role in GSC maintenance, we observed that germline cells, including GSCs, were almost completely absent from the atrophied ovaries from *p53* transheterozygous dysgenic females (Figure 4A). By contrast, *p53* heterozygotes frequently exhibited developing oocytes at all stages, including GSCs (Figure 4C).

Although we observed no evidence for normal oogenesis at any stage in the atrophied ovaries of *p53* transheterozygous dysgenic females, we did encounter a number of isolated cells the express the germline-specific marker Vasa (Figure 4B). However, these cells did not exhibit the spectrosome structures that are characteristic of GSCs or CBs, nor were they found in multicellular cysts that characterize subsequent stages of oogenesis. The origin of these germline cells is unclear, but they suggest a disruption in the early oogenesis, in which stem cell identity is lost, but mitotic divisions and differentiation do not occur.

**Figure 4.**
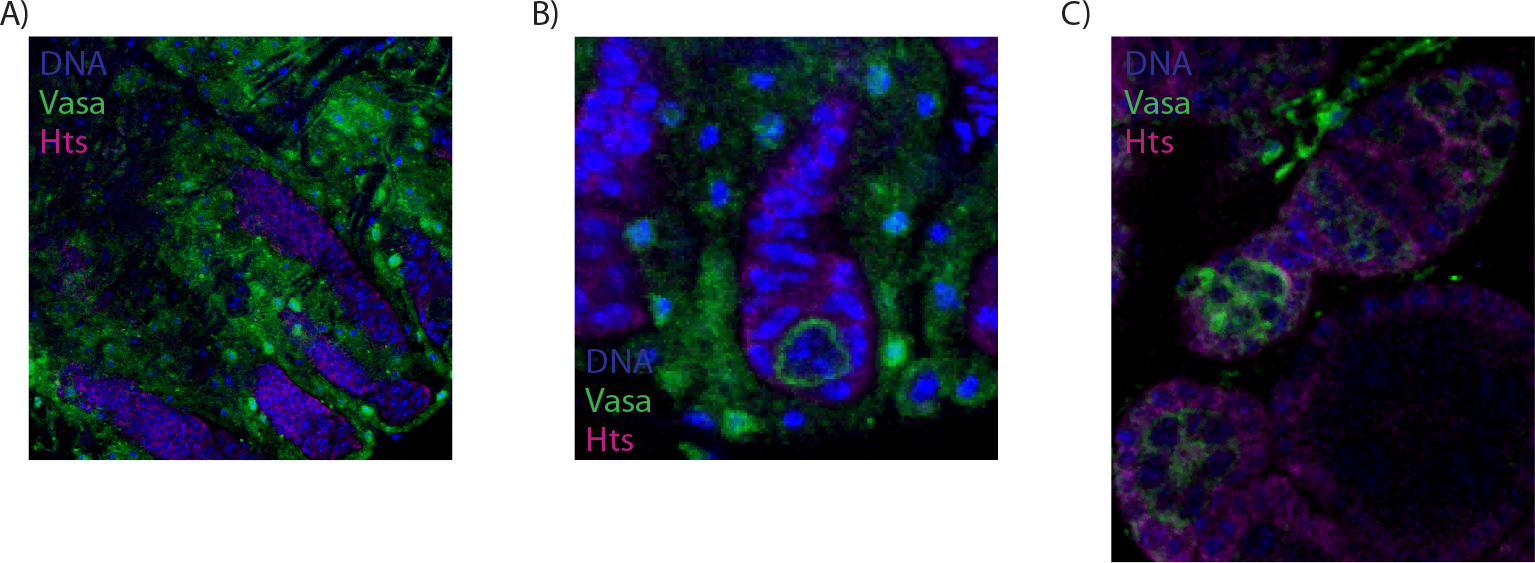
Atrophied ovaries of *p53* mutant dysgenic females lack GSCs. Germaria from transheterozygous and heterozygous *p53* dysgenic females. Vasa, which is found in germline cells and the ovarian sheath, is shown in green. Hts-1B1, which is found in circular spectrosomes in GSCs and CBs, as well as somatic cells that surround the developing egg chamber, is shown in magenta. DAPI-stained DNA is shown in blue. A) Atrophied ovaries from transheterozygous *p53* dysgenic females contain empty ovarioles that lack Vasa-positive germline cells. B) Occasional isolated germline cells, lacking spectrosomes, are found in *p53* transheterozygous dysgenic ovaries. C) Ovaries of *p53* heterozygotes often exhibit GSCs and CBs in germaria, as well as developing egg chambers.

## CONCLUSION

Transposable elements impose a significant burden on their host genome, not only as a cause of deleterious mutations (Dupuy et al., 2001; Spradling et al., 1999), but also as a source of genotoxic stress (reviewed in Castañeda et al., 2011; Hedges and Deininger, 2007). As arbiters of cellular response to DNA damage, the DDR plays an obvious yet poorly understood role in the determining how host cells are affected by TE activity. Using *P*-element hybrid dysgenesis as a model, our observations reveal that DDR components Chk2 and p53 are determinant of cell-fate in the presence of unregulated transposition. Our observations further highlight the unique role of p53 in germline stem cells in conferring tolerance of transpositional activity, by preserving GSCs in the face of increased DNA damage.

The molecular function of p53 protein in *P*-element dysgenic germlines remains unclear. However, p53 was recently revealed to act as a negative regulator of retrotransposon activity in the female germline (Tiwari et al., 2017; Wylie et al., 2016). Interestingly, TE derepression in *p53* mutants depends on the meiotic nuclease *spo11*, suggesting that retrotransposon activation depends on the production of double-stranded breaks (Wylie et al., 2016). *P*-element transposition in GSCs and CBs may therefore act as a trigger, which, in the absence of *p53* releases regulation of retrotransposons, causing enhanced DNA damage and germline loss.

## ACKNOWLEDGEMENTS

We are grateful to John Abrams, Haifan Lin, Richard Meisel, Jeff Sekelsky and William Theurkauf for providing mutant and transgenic *Drosophila.* We are also grateful to members of the Kelleher and Meisel laboratories for helpful discussion. Sadia Tasnim was supported by a Summer Undergraduate Research Fellowship from the University of Houston. Erin Kelleher was supported by NSF DEB #1457800.

